# Pro-neurogenic potential of blood cells cultured in autologous plasma

**DOI:** 10.1101/2025.03.19.643269

**Authors:** Rhythm Arora, Alka Bhardwaj, Naresh K Panda, Reena Das, Sanhita Sinharay, Pooja Patkulkar, Tulika Gupta, Jaimanti Bakshi, Ramandeep Singh Virk, Gyanaranjan Nayak, Sourabh K Patro, Sanjay Kumar Bhadada, Arnab Pal, Rimesh Pal, Seema Chhabara, Sanjeev Chhabra, Meenakshi Pal, Sajid Rashid, Maryada Sharma

## Abstract

The lack of effective stem cell protocols for generating personalized neurovascular niches poses a critical challenge in precision medicine. While iPSC-based methods are explored, their clinical use is hindered by high costs, long timelines, and cancer risks. Recent advancements in plasma-driven differentiation, using circulating monocytes, offer a promising solution as they can be reprogrammed into neuron-like, endothelial-like, and hematopoietic cells without genetic manipulation, by inducing growth factors mediated transdifferentiation. Vasculature is integral to neurodevelopment, with early blood supply transitioning from the perineural to intrinsic vascular plexus, driven by neuro-hematovascular signaling. The choroid plexus selectively transports proteins and growth factors from blood to CSF, supporting neural proliferation and differentiation. Building on these insights, we leveraged the innate reprograming potential of blood-derived cells to generate neuro-hematovascular niches using a novel PITTRep methodology, devoid of transgene and growth factor mediated transdifferentiation opening new avenues for regenerative and investigative neurovascular studies.

## Introduction

Paucity of protocols that can co-induce individualized neurovascular niches is an unmet challenge. Therefore, it is envisioned that the generation of *in vitro* vascularized neuronal niches from easily accessible donor source cells could represent a potential framework to investigate the altered neurovascular pathways in precision medicine targeted investigative studies. The aspect of using pluripotent stem cells and inducing reprogramming by transgene overexpression or morphogenetic factors in special culture media is cost ineffective. Further, the most concerning aspect is potential presence of cancer-driving mutations in induced pluripotent cells that dissuades their use in clinical application (Lezmi et al., 2024). Generation of transgene-free neuro-hematovascular cells from PSCs is encouraged for clinical applications. Reprograming protocols that employ serum-free media, and are driven by developmental principles have encouraged establishment of hematovascular (vessel-like cells lining hematopoietic clusters) cell types following differentiation of spinning embryoid bodies into hemogenic endothelium lineage intermediates or selective expression of engraftable hematopoietic niches (Sroczynska et al., 2009; Ng et al., 2016; Ng et al., 2024) A single-stage human plasma-dependent differentiation of embryoid bodies was also reported to result in engraftable hematopoietic stem cells (Piau et al., 2023). A subpopulation of circulating monocytes is demonstrated to be amenable to pluripotency induction by growth factors, followed by non-genetic transdifferentiation into phagocytic cell types, endothelial-like cells, and neuronal-like cells (Bellon et al., 2018; Zhao et al.; Kuwana et al., 2003; Horschitz et al., 2010; Ruhnke et al., 2005; Kodama et al., 2006; Liu et al., 2011; Bellon, 2024). Importantly, neural stem cells are reported to differentiate into a variety of hematopoietic cells, and circulating lymphoid cells and macrophages are known to enter the brain (Knopf et al., 1998; Williams and Hickey; Hickey, 1999), where migrating bone marrow cells can differentiate into microglia and astrocytes following transplantation into irradiated recipients (Theele and Streit, 1993; Eglitis et al., 1997). Also, tissue-cultured bone marrow stromal cells display neuronal markers (Sanchez-Ramos et al., 2000; Woodbury et al., 2000), and upon transplantation into lateral ventricles were shown to differentiate into astrocytes (Azizi et al., 1998; Kopen et al., 1999). Therefore, it was suggested that a continuous influx of bone marrow stem cells into the ependymal and subependymal zones can contribute to generation of various CNS cell types. Exposure to the CNS environment was suggested to play an important role in induction of neuronal phenotypes in bone marrow derived mesodermal lineage cells that were found to home to cortex, hippocampus and other complex areas of the brain. Bone marrow is relatively more accessible as opposed to brain’s ventricular or sub granular neural stem cells, and they come with an advantage of having inherent host compatibility; therefore, they have been proposed to be promising sources of alternative neural stem cells (mezey et al.,2000; Brazelton et al.,2000).

The above studies reflect the inherent potential of blood-derived cells for neuronal developmental reprograming, and pro-hematopoietic differentiation potential of plasma components. The current literature is replete with several protocols that are based on vascularized brain organoids, assembloids, and building neurovascular unit (Cakir et al., 2019 Rizzuti et al., 2024; Sandoval et al.,2024. However, to the best of our knowledge the potential co-induction of neuro-hematovascular niches from autologous blood components and their attendant homeostasis protease cascade (central to wound healing) has never been explored till date. Based on preliminary observations from our lab that suggested upregulation of neuro-immune pathways in plasma-cultured monocytic cells (Sharma et al., 2022) we tested the possibility of induction of neuro-hematovascular niches in leukocytes cultured with autologous plasma (Arora et al., 2024, **Patent Application no.202411034351)** We report successful establishment and characterization of neuro-hematovascular niches by employing original method that combines prolonged culturing of autologous blood components (leukocytes and RBCs cultured in plasma) for induction of neuro-hematovascular niches, termed as PITTRep (<u>P</u>lasma Induced epi<u>T</u>ranscriptomic and <u>T</u>ranscriptomic <u>Rep</u>rograming). The emerging bipotential neuro-hematovascular progenitor colonies are termed as haemGOD bodies (haem/blood-derived Glial Ontogenic Determinants) that exhibit de novo patterning into neurovascular niches when cultured in autologous plasma.

## Results

### De novo induction of neuro-hematovascular niches from autologous blood

Previous studies have tried to generate hematopoietic stem cells and neurons from peripheral blood mononuclear cells but coexistence of these niches together in a system has never been reported. Recent study by the Elizabeth and group had used a small fraction of plasma along with other growth factors to develop hematopoietic stem cells from induced pluripotent stem cells (Ng et al., 2024). This inspired us to use autologous blood components i.e. leukocytes, erythrocytes and plasma to develop a model which exhibits co existence of ectodermal lineage neuronal cells and mesodermal hematopoietic and vascular cells.

The blood cells were isolated using gradient dependent centrifugation and then mixed with autologous plasma and plated in culture plate with around 3.5-4.5 million WBC’s /ml plasma (the cell count varies donor to donor) (Figure 1A). On day1, we observed aggregation of cellular clusters with golden brown margins, in our culture which increase in size and number as the culture progresses (Figure 1Bii and iii). We named them hemeGOD bodies (Heame(blood) derived Glial Ontogenic Determinants) the reason for which will be elaborated further. These hemeGOD bodies begin to radiate at around day 7 (Figure 1Bv), and flattened elongated like cells emerge out of them resembling the endothelial and radial glia morphology. Interestingly, we noticed a small population of shiny pearl like appearance cells sprouting from these bodies, of the diameter 6 *µ*m, this population resembles the diameter of hematopoietic stem cells (Figure 1Ci and ii). As the culture progresses the sprouting of the hemeGOD bodies increases and by day 10 the culture is full of elongated cells and hematopoietic stem cell like population. By day 14, extensive tissue formation takes place and the appearance of these cell types relatively decreases (Figure 1Bix). These findings were obtained from the upper phase of the culture, were the cells are freely suspended in the plasma and ultimately lead to tissue like aggregation. During routine microscopic observation when focused on the surface of the well in the culture plate we noticed the development of faint fine needle like projections in the culture plate, here on referred as lower phase around day 3 (Figure 1Bx). These projections became more noticeable as the culture progresses. By day 7, there number increased and exhibited increased in neurite like processes (Figure 1Bxi). At day 10 we noticed, rigorous increase in the network pattern which seemed to be developing neurons as their morphology possessed fine processes resembling dendrites (Figure 1Bxii). By day 14 the network is less visible as the culture rapidly forms a tissue like aggregation and the lower phase can’t be focused precisely.

**Figure 1:**
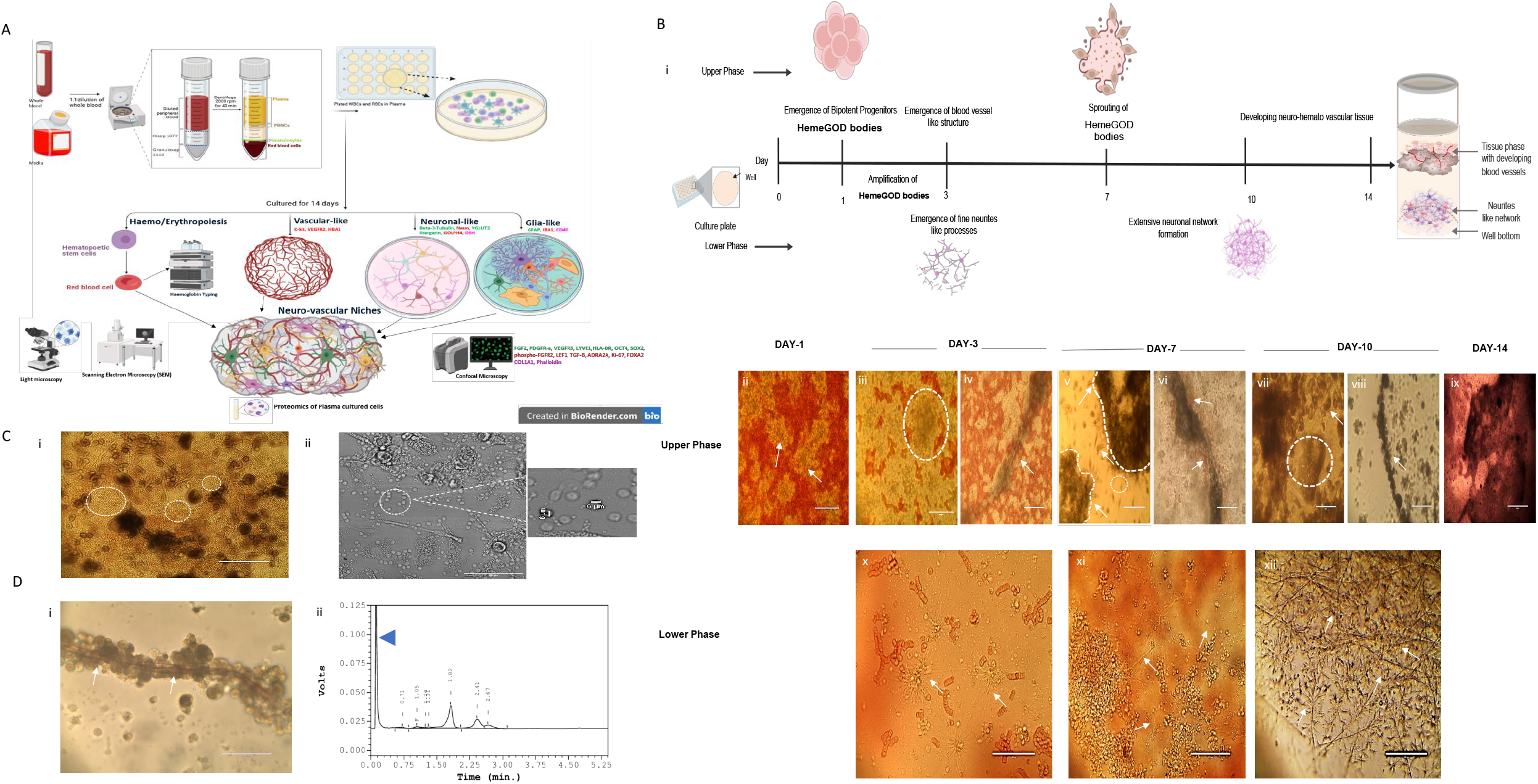
De novo induction of neuro-hemtaovascular niches from autologous blood A) Schematic showing detailed methodology of culturing neuro-hematovascular niches from autologous blood components like leukocytes, erythrocytes and plasma without any transgene approach with details of various tools and techniques used to characterize them B) (i) Schematic illustration showing culture progression of the developing neuro-hematovascular tissue growing in autologous plasma showing key highlights of the culture in both the upper and lower phase. (ii) White arrows indicate the emergence of golden brown cell clusters at day1 named as heameGOD bodies. (iii) HeameGOD, marked in white dotted circle increase in size by day 3. (iv) Blood vessel like structure observed at day 3. (v) Sprouting of the HeameGOD bodies (marked in dotted white line) and release of flattened elongated cells indicated with white arrows and small shiny pearl like appearing cells indicated in white dotted circle. (vi) Presence of newly emerged erythrocytes around the blood vessel indicated with arrows. (vii) Increase in the number of flattened elongated cells (white arrow) and the shiny pearl like cells emerging from heameGOD bodies (white dotted circle). (viii) Clustering of cells (white arrow) increases around the blood vessel. (ix) Formation of tissue like structure in the well. (x) Fine neurite like processes emerging at day 3 (xi) and (xii) Extensive development of the neuronal network as observed in the lower phase. C) (i) Light microscope captured image of Day 10 culture showing rigorous formation of shiny pearl like cells. (ii) On measuring the diameter of these cells in brightfield captured via confocal microscope it turns out to be 6μm. D) (i) Presence of erythrocytes in the developing blood vessel as seen on day 14. (ii) Hemoglobin typing indicating the formation of peak corresponding to development hemobglobin appearing between 0.00-0.05 mins. Scale Bars-B(ii, iii, vii, ix,x, xi, xii), C(ii) and D(i)-100 μm B(iv, vi, viii) and C(i)-200 μm

Notably, we saw the emergence of blood vessel like structures at day 3 which grew in size and got surrounded by hemeGOD bodies over the period (Figure 1 Biv, vi, vii). We noticed the lumen of blood vessel being filled with blood around 14 day (Figure 1Di). Which made us question if erythropoiesis was taking place in our culture. To confirm this we performed hemoglobin typing of culture (Figure 1Dii) and the results indicated the peaks of developmental Hb in our system which could otherwise be not found from adult hemoglobin that we add during culture.

It is important to understand that, no culture milestones appear as single stage at any day, when we specify about a culture milestone, we mention it in relative context to the other components. The system is completely physiologic and shows de novo growth of heterogenous cell types making them tough to identify from the morphological characteristics. The culture is not regulated by any artificial supplementation hence, we do not select any specific cell type proliferation.

In order to validate our microscopic findings, we performed immunofluorescence on our cultured cells at various time points. We tested our culture for pluripotency markers such as Oct4 with marker of cellular proliferation Ki67 (Figure 2Ai). The hemeGOD bodies showed strong positivity for both these markers which suggests that they possess stemness and are actively dividing. We hypothesized that these hemeGOD bodies are the bipotential progenitors which are giving rise to cells of ectodermal and mesodermal lineages hence we tested them with SOX2, a neural progenitor marker and ADRA2A, a marker for adrenergic control of hematovascular niches, the sprouting cells from hemeGOD bodies show immunoreactivity to ADRA2A but no SOX2 positivity (Figure 2Aii.). We further validated our hypothesis of bipotential nature of haemGOD bodies by testing these bodies with GFAP, expressed in glia progenitors and c-Kit or CD117 a maker of hematopoietic progenitor cells, to ensure there bipotential nature, and the sprouting hemeGOD bodies showed positive expression for both the makers (Figure 2Aiii).

**Figure 2:**
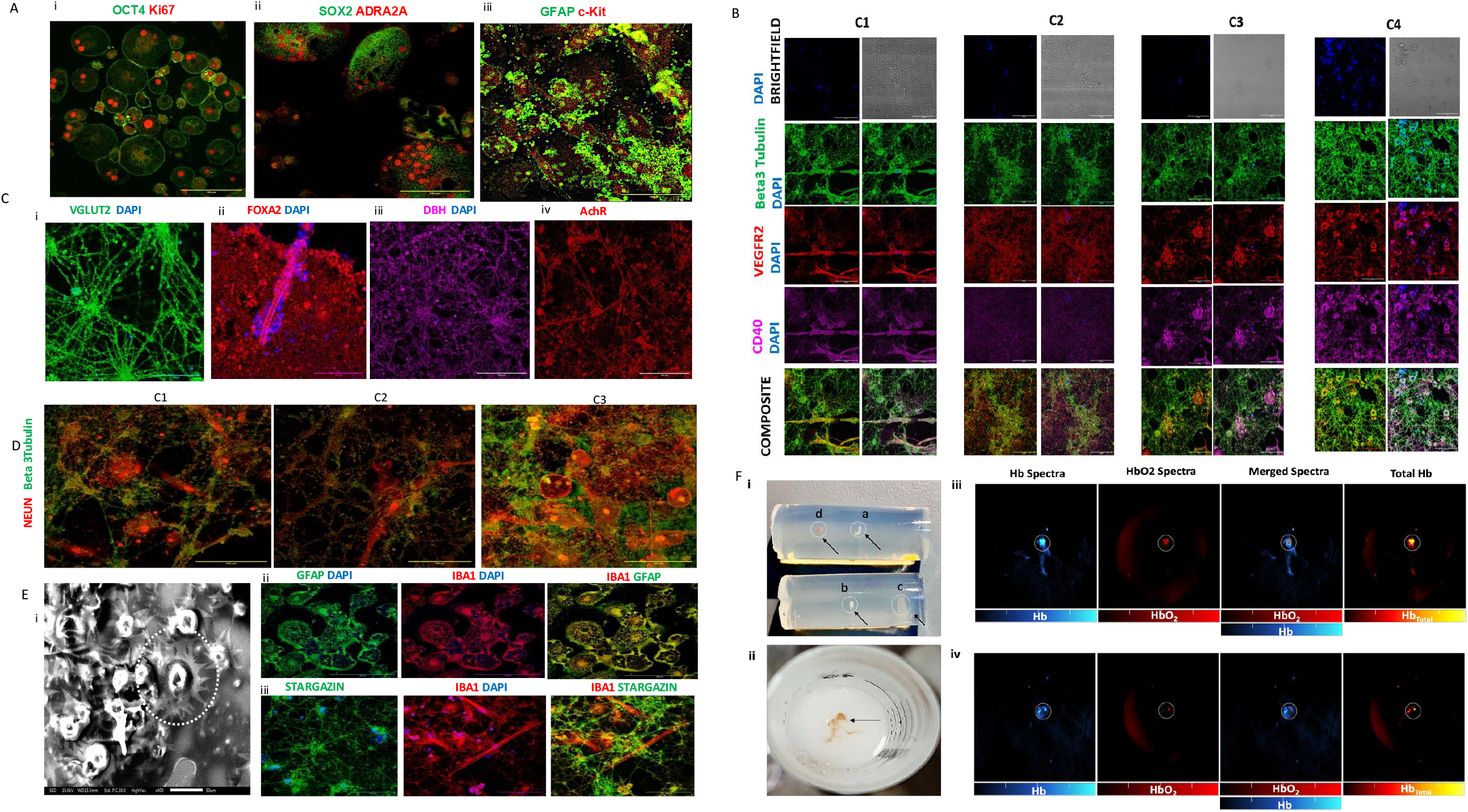
Characterization of neurohemtaovascular niches A) (i) HeameGOD bodies express pluripotency marker Oct4 and Ki67 cellular proliferation marker. (ii) Sox2 expression in sprouting heameGOD bodies (iii) GFAP positive cells sprouting from C-kit positive heameGOD bodies indicating there bipotential nature. B) Establishment of a neuro-hematovascular niche in day 14 culture, the culture shows strong positivity for Beta-3 tubulin, expressed in neurons, CD40 and VEGFR2 expressed in endothelial cells. The co-expression of these markers increases when we increased the volume of RBC of in culture from C1 to C4. Where C2 serves as baseline added volume of RBC, C3 has twice the volume and C4 has three times the volume than C2 and no additional RBC were added in C1. C) Immunostaining for identifying different types of neurons in culture. (i)VGLUT2 (Vesicular glutamate transporter-2)for glutamatergic neurons(ii) FoxA2, a transcription factor for promoting midbrain dopaminic neurons ((iii) DBH (Dopamine beta-hydroxylase) for adrenergic and non-adrenergic neurons (iv) AchR (Acetylcholine receptor) for cholinergic neurons. Nuceli in C(i) – C(iii) are stained in DAPI. D) The expression of neuronal maturation marker increases as we move from C1-C3 as compared to Beta-3 tubulin, expressed by early neurons. E) (i)Scanning electron microscopy image capturing glia cell morphology with ramified process (marked in white dotted circle) at day 14. (ii) Iba-1 positive heameGOD bodies showing strong positive GFAP expression (iii) Iba-1 expressing microglia (marked with white triangles), present in association with Stargazin positive neurons. Scale bars A (i-iii), B, C(i-iv), D, E (ii-iii) 100 μm, E(i)50μm F) Optoacoustic imaging indicating the formation of neuro vascularised tissue: i) Neurovascularized tissue (indicated by black arrows) of the conditions C1-C4 embedded in cylindrical phantoms prepared from low melting agarose where a= C1 condition, b = C2 condition, c= C3 condition and d= C4 condition of the cultured tissue. ii) A closer view of iv (C4 condition) neuro vascularised tissue placed in low melting agarose for optoacoustic imaging. iii) and iv) are scans of the neuro vascularised tissue (d and c respectively) obtained from the iTHERA MSOT inVision optoacoustic imaging device. Notice the defined Hb and HbO2 signal in localised in the boundary of neuro vascularised tissue. Some portions of this neuro vascularised tissue are richer in signal assigned to Hb and HbO_2_ (highlighted in circle) than the other parts. There is more intensified Hb-HbO2 signal in the scan of C4 condition (iii) than C3 condition (iv) neuro vascularised tissue.

Since we observed both the expression of vascular and neuronal progenitor cell types in the same culture, we tested our 14^th^ day culture with Beta 3 tubulin, exclusively expressed in neurons along with vascular/immune markers, CD40 and VEGFR2.The culture showed strong positivity for these markers, and their expression increases as the concentration of RBC’s increased indicating that increasing concentration of RBC facilitates rigorous development of neuronal and vascular niches (Figure 2B). The strong positive expression of beta 3 tubulin confirmed the presence of developing neurons in our system. These findings intrigued us to test for the types of neurons being developed in our model. Thus, we performed immunostaining with different markers like VGLUT2, expressed in glutamatergic neurons, Foxa2, which regulates dopamine neuron generation and differentiation DBH, expressed in adrenergic and noradrenergic neurons and AchR, expressed in cholinergic neuron receptors (Figure 2Ci-iv). The culture showed strong immunoreactivity for all these markers strongly suggesting the de novo development of various types of neurons in the culture. Staining with Neun, a neuronal maturation marker, suggested that with increase in the RBC concentration the tissue shows increased expression of neuronal maturation marker (Figure 2D).

Astrocytes and microglia form the essential supporting niche for neuronal maturation and synapse formation. Although both the cell types arise from different progenitors i.e. astrocytes differentiate from neuronal progenitors and microglia originate from embryonic hematopoietic precursors we suspected if the bipotent hemeGOD bodies could be the precursors for these cell types. Despite the fact we could not directly differentiate between the two cell types in brightfield microscopy, because of their similar morphology, results from scanning electron microscopy revealed the presence of small body ramified process like cell type in our culture similar to glia like morphology (Figure 2Ei). In order to confirm if these cells were microglia or astrocytes, we tested our culture with Iba-1 a classical microglia marker and GFAP expressed by astroglia progenitors (Figure 2Eii). The hemeGOD bodies have taken up both the stains supporting there bipotential nature and suggesting that both microglia and astroglia progenitors originate from a common precusor in our system. Immuno staining with Iba-1 and Stargazin, which regulates neuronal excitability and plays a crucial role in neuronal plasticity in glutamatergic neurons shows presence of microglia cell type in our system (Figure 1Eii).

To further test the development of functional vascular components in our growing cuture, we performed some preliminary experiments by imaging the live neurovascular tissue using a multispectral optoacoustic tomography scanner (MSOT, iTHERA Medical™, Germany) (Figure 2 F). The tissue was examined by preparing a phantom (Figure 2 F i and ii) and optoacoustic signals were acquired for the wavelength range of 660-980 nm with 5 nm intervals. This long wavelength range was selected to ensure the characterization of any chromophores in vascularized tissue that absorb in the near infrared wavelength range (NIR). Post processing and spectral unmixing of oxygenated and deoxygenated hemoglobin signatures from the acquired images indicated the presence of a defined Hemoglobin (HbO2) and Deoxy-hemoglobin (Hb) signal in the neuro-vascularised tissue (Figure 2iii). Although no micro vessels got defined from the Hb and HbO2 spectra scans. The detection of Hb and deoxy-Hb spectra in optoacoustic imaging confirms the presence of functional vascular components within the neurovascular tissue. This indicates the ability of the tissue to support oxygen delivery and consumption, key features for mimicking physiological neurovascular function and studying brain-related pathologies.

### Plasma supports growth factor signaling of neuro-hematovacsular niches

The platelet containing, plasma obtained from density dependent centrifugation, is rich in growth factors like fibroblast growth factor (FGF), transforming growth factor-beta (TGF-β), insulin like factor, vascular endothelial growth factor (VEGF), platelet derived growth factors (PDGF) and other free circulating growth factors. During blood collection, there occurs a mild activation of platelets which contributes to the pool of growth factors. We hypothesized that plasma growth factors promoted growth factor signaling in our model which helps in culturing multilineage cell types neurons, glia, endothelial and hematopoietic stem cells and promotes vasculogenesis. For testing our hypothesis we tested our culture with heterogenous cell types with various cell signaling receptors. FGF-FGR2 signaling, which promotes neurogenesis (Woodbury and Ikezu, 2014), angiogenesis (Jia et al., 2021,Murakami and Simons, 2008) and stem cell proliferation (Mossahebi-Mohammadi et al., 2020). FGF^+^/PhosphoFGFR2^+^ expression indicates active FGF signaling occurring in our system(Figure 3Ai and ii). PDGFRα, platelet derived growth factor receptor alpha expression in culture is suggestive of active PDGF signaling pathway supported by platelet-derived PDGF in the plasma (Figure 3Bi and ii). Strong expression of VEGFR2 and VEGFR3 aligns with the presence of VEGF signaling crucial for active vascular differentiation signaling as seen in our culture(Figure 3B and C). The culture shows strong positivity for TGF-β^+^ essential for neurogenesis and gliogenesis (Meyers and Kessler, 2017) which explains the development of these cell types in our model(Figure 3Ci and ii). TGF-β often regulates collagen deposition necessary for maintaining structural integrity of the developing tissue (Deng et al., 2024). We analyzed the formation of collagen in the developing system which was confirmed by the positive expression of COL1A1(Figure 3Ci and ii). Canonical Wnt signaling activated transcription factor LEF1 was also found in our system suggestive of active Wnt signaling(Figure 3D). IGF levels were calculated using IGF-1 quantitative assay from the cultured plasma and levels were compared between the cultures with low RBC concentration which shows more cellular stages and high RBC concentration which promotes more tissue formation, the results show significantly high expression of IGF in tissue stages, almost twice as present in cellular stages. This highlights that IGF promotes tissue formation (Figure 3F). GFAP positive structures are seen interacting with TGF-β^+^ cells (Figure 3E).

**Figure 3.**
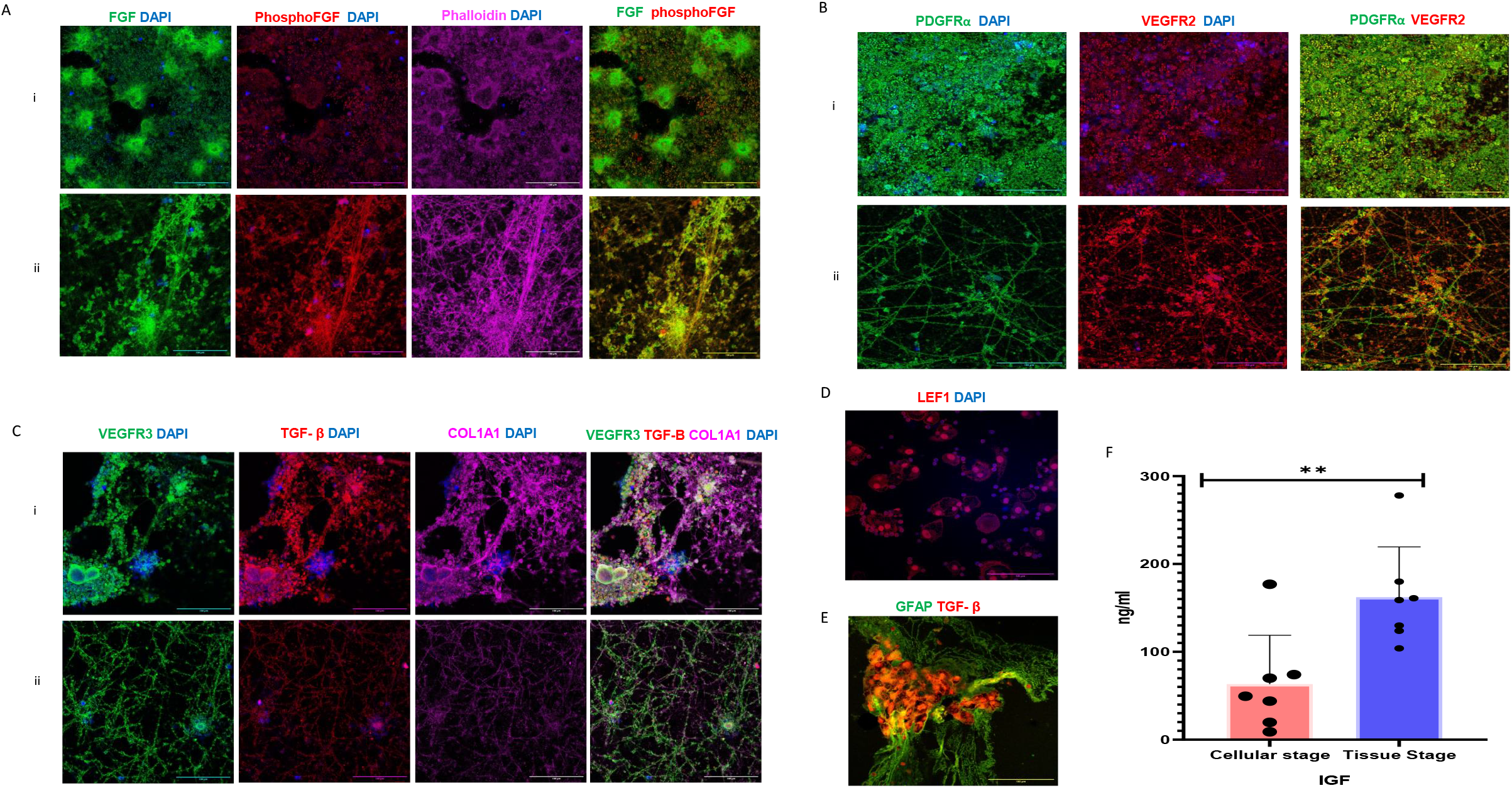
Plasma supports growth factor signaling of neuro-hematovacsular niches A) (i) and (ii) FGF(Fibroblast growth factor) and PhsophoFGFR2(Phosphorylated fibroblast growth factor receptor 2) expression across the heterogenous cell types in the growing neuro-hemtaovascualr niches at day 14 indicative of active FGF signaling. Phallodin is used for staining actin and nuclei are stained with DAPI. B) (i) and (ii) PDGFRα (Platelet-derived growth factor receptor alpha) and VEGFR2 (Vascular Endothelial Growth Factor Receptor 2) expression across the heterogenous cell types in the growing neuro-hemtaovascular niches at day 14. C) (i) and (ii) Heterogeneous expression of VEGFR3 (Vascular Endothelial Growth Factor Receptor 3) and TGF-β (Transforming growth factor-beta) at day 14 of culture. COL1A1 (Collagen type I alpha) is used to stain collagen in developing tissue. D) LEF1 expression at day 14 in the cutlre. DAPI is used to stain the nuclei. E) GFAP positive cells emerging from TGF-β positive cells. F) Graph showing significant increase the concentration of IGF (Insulin-like growth factor) in tissue stages of neuro-hematovascular niches when compared cellular stages. Scale Bars: A(i,ii), B(i,ii), C(i,ii), D, E-100μm

For a heterogeneous culture like ours it tough to comment which cell type is responsible for which growth factor signaling. Further experiments are being carried to distinguish the key cellular players involved in the cross talk for developing this dynamic system. Expression of various growth factors and growth factor receptors in our model elucidates a crucial role of growth factor rich plasma which acts as a natural biomimetic microenvironment for the delicate molecular cross talk to take place. This depicts plasma as a strong universal medium which supports differentiation and maintenance of multiple lineages like in our system showcasing its potential in translational and clinical applications.

### Plasma mediated complement activation drives induction of neuro-hematovascular niches

By now we have been able to establish that plasma mediated growth factors promote growth factor signaling for developing neuronal, glial, endothelial and hematopoietic stem cell niches from differentiated leukocytes. In order to understand how plasma is able to induce the adult blood cells of mesodermal origin to neurons which are ectodermal in origin and multipotent stem cells like hematopoietic stem cells we performed LCMS/MS on our cultured cells and compared it with cells cultured in RPMI with 10% FBS under routine culture conditions. The proteomics analysis revealed a significant upregulation of blood coagulation and complement pathways genes in cells cultured in plasma compared to the control (Figure 4A and B). This activation, observed in plasma-cultured cells, signifies the interplay between intrinsic and extrinsic pathways, further amplified by increased thrombin generation (Pryzdial et al., 2022; Luo et al., 2020, Wiegner et al., 2016, Hamad et al., 2008, Markiewski et al., 2007). Plasmin plays a crucial role in regulating the homeostasis of the physiologic complement coagulation cascade (Pryzdial et al., 2022; Luo et al., 2020, Wiegner et al., 2016, Hamad et al., 2008,Strande and Phillips, 2009). In order to validate the activation of blood coagulation complement cascade and the role of RBC’s in mediating neuro-hematovascular tissue niches we performed functional thrombin generation and plasmin activity assays in our culture conditions from C1-C4 which shows a significant increase for both thrombin and plasmin activity (Figure 4 E and F).

**Figure 4.**
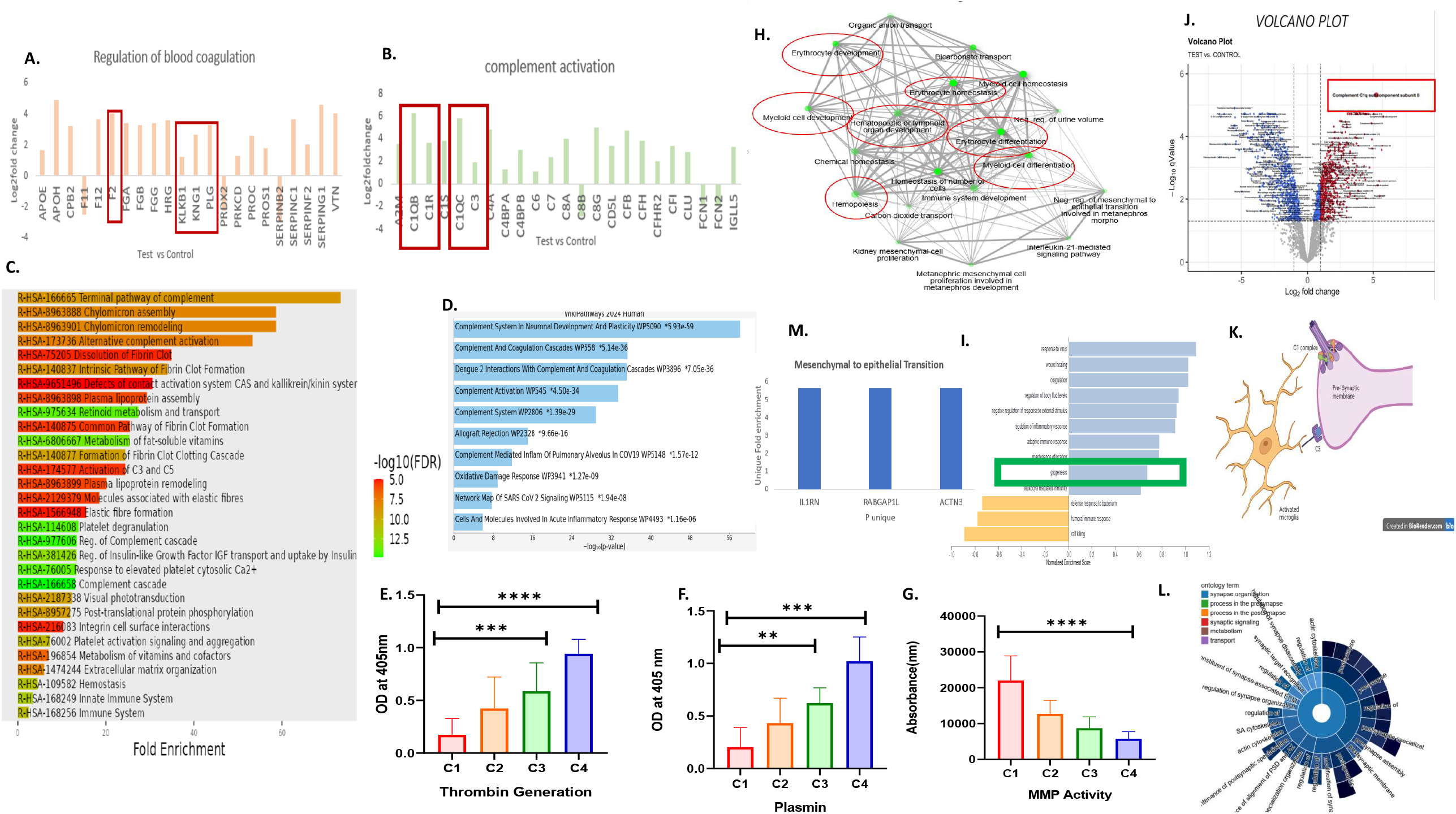
Comprehensive Bioinformatic analysis and Quantitative Validation of Proteomics Data from Plasma-Cultured Cells (A-B) Bar plot showcasing upregulated blood coagulation proteins, indicating activation of intrinsic and RPMI and 10%FBS cultured cells, particularly C3 and C1q components. (C) A Reactome enrichment plot created on shiny GO v0.80, highlights key pathways, including complement-coagulation, IGF signaling, and extracellular matrix (ECM) organization. (D) Wiki-Pathways analysis of highly significant proteins, points to important roles of complement in neuronal development and synaptic plasticity. (E-G) Functional assays validate key enzymes of complement-coagulation cascade, thrombin generation, increasing with increased concentration of RBCs(E), plasmin activity also increases with increased concentration of RBCs (F), and matrix metalloproteinase (MMP) activity decreasing with increase concentration of RBCs (G) (H) A pathway-pathway interaction network(Shiny GO v 0.80) created using significant proteins in plasma cultured cells revealing processes like erythropoiesis and hematopoiesis. (I) Pathways associated with gliogenesis are enriched, emphasizing the roles of gliogenesis and complement in neurogenesis. (J) A volcano plot created on Cytoscape indicates significant upregulation of complement component C1qB in plasma cultures. (K) A schematic illustrates complement-mediated neuronal pruning involving C1q and C3. (L) SynGO 2022 plot showing Synaptic cytoskeleton organization indicative of active regulation in plasma cultured cells. (M) Bar plot showing unique proteins expression in plasma cultured cells leading to mesenchymal to epithelial transition.

Further analysis using Shiny GO v0.80, Enrichr, and Appyter demonstrated that plasmin activation is proportional to thrombin generation, activating complement components C3 and C5 (Pryzdial et al., 2022; Luo et al., 2020, Wiegner et al., 2016, Hamad et al., 2008, Markiewski et al., 2007,Choi et al., 2018). The detailed pathway analysis revealed the activation of complement factors C3 and C5, followed by platelet activation and degranulation, as well as the regulation of insulin growth factors essential for neurogenesis. Supporting data from Wiki-Pathways 2024 (Figure 4 D) indicated a significant pathway of the complement system in neuronal development and plasticity which suggests role of complement coagulation in neurogenesis in the plasma cultured blood cells.

We speculate a possibility, that activation of growth factors through plasmin and thrombin, following stimulation of C3, C5, and C1q components of the complement cascade, may contribute to the direct trans-differentiation of blood cells cultured in plasma into neuronal cell population. To establish a possible mechanistic basis of this intriguinging process, we did comprehensive bioinformatic analysis of day 4 proteomics data of plasma cultured cells and found that a combination of unique genes (IL1RN, RABGAP1L, ACTN3) were linked to the upregulation of two important pathways, mesenchymal-to-ectodermal transition (MET) pathway and gliogenesis (Figure 4I and M). We propose that these common genes are the crucial players in regulating the induction of blood cells to ectodermal glial progenitors in the culture. These blood-derived progenitors with glial ontogeny are unique to the PITTRep reprograming method as they give rise to mesodermal-origin microglia and ectodermal origin neuronal progenitors. As indicated in Figure 2E ii by the strong immunoreactivity of Iba^+^/GFAP^+^ cells emerging from the common hemeGOD bodies. Since these cKIT^+^/GFAP^+^ glial progenitors (Figure 2A iii) are emerging from mesodermal blood cells hence the name Heme-derived Glia Ontogenic Determinants was given to this population.

Volcano plot created using Cytoscape (Figure 4J) identified C1qB as the top upregulated protein in our plasma cultured cells. Complement mediated synaptic pruning regulated via the C3 and C1q components of the complement cascade is an important phenomena observed in microglia cells is essential for building strong neuronal connections(Hilton et al., 2024, Lindhout et al., 2024, Pereira-Iglesias et al., 2025, Kojima et al., 2010 Matsuda et al., 2013, Hosokawa and Liu, 2021, Fadahunsi et al., 2024) which promotes post synaptic organization (Figure 4K and L) can be seen in our proteomics data**Stargazin**^**+**^**/Iba**^**+**^ **immunostaining reveals the presence of this phenomena in our culture (Figure 2 Eiii)**. The maintenance and regulation of physiological neurovascular niches requires proper extracellular matrix (ECM) organization, is a significant upregulated pathway in the proteomics analysis (Figure 4C) (Long and Huttner, 2022, Wallace and Pollen, 2024, Pillai and Franze, 2024). Results from immunostaining also reveals the strong expression of COL1A1 in the developing neuro-hemtaovascular niche (Figure 3Ci and ii). Matrix metalloproteinases (MMPs) which play a crucial role in regulating the proteolytic degradation of ECM essential for tissue remodeling and development, are found to have descending activity trend with the increasing tissue formation (Figure 4G). Detailed proteomics analysis revealed the development and differentiation of myeloid cells, erythrocyte development, and hematopoiesis (Figure 4H), supporting the hemoglobin typing data (Figure 1Dii).

## Discussion

Limited access to live human brain tissue encourages establishment of a structural and functional neurovascular unit (a complex multi-cellular system consisting of endothelial cells, neurons, glia, smooth muscle cells, and pericytes) to comprehensively understand the sensory-evoked brain activity mechanisms underlying neurosensory development and deregulation. However, faithful *in vitro* models with robust neurovascular niches that exhibit phenomenon of neurovascular coupling are lacking. Therefore, establishment and characterization of donor-derived neuro-hematovascular niches is a prerequisite to understand the developmental and pathological trajectories linked to neurosensory and neurodevelopmental disorders. Brain organoids have remarkably transformed our understanding of early brain development by reflecting the species-specific developmental mechanisms and informing on cellular phenotypes linked to human brain diseases; however, they have key limitations that restrict their use in understanding brain-activity related mechanisms. Physiological and pathological neurological pathways are guided by overlapping morphogenic cues, that encourages the use of self-patterning approaches over exogenous morphogens to drive organoid development. Integration of sensory inputs by generating brain and inner ear organoids/retinal organoids is proposed to simulate sensory-evoked physiological patterns in brain (Birtele et al., 2024). However, establishment of successful neurovascular niches with active blood flow is a prerequisite to map functional brain-activity patterns that is currently possible only through clinical functional neuroimaging techniques, which are based on magnetic and optical properties of hemoglobin, making presence of erythroid and vascular lineage cells indispensable for functional mapping of brain.

From an evolutionary aspect of brain development, the vasculature is emerging as a key contributor to brain morphogenesis and function during neurodevelopment. Neurogenesis, astrogliogenesis, and oligodendro-gliogenesis overlap with CNS vasculature, and choroid plexus development. Interestingly, neural progenitors (subventricular zone, subgranular zone, and transient rhombomeres) in brain unequivocally arise close to ventricular areas enclosing CSF, that forms an important (perhaps indispensable) microenvironment for the growth and proliferation of these progenitors. The hydrodynamic properties of CSF and the relative high blood flow rate of choroid plexus microvasculature compared to other brain areas (cortex with relatively low blood flow rate) emphasize the importance of the plexuses for exchange across this interface early in development, and these active hydrodynamic properties may contribute to enriched ventricular apposed neural progenitor zones as opposed to distant differentiated neuronal populations. The current study harnessed the (potential yet unexplored) hemorheological properties of RBCs in context of neuro-hematovascular development. The nature of blood flow is non-Newtonian and is a function of RBC concentration /haematocrit when they flow through the regions of low shear rates as seen in cerebral microcirculation. The current reprograming approach is a modification of our previous cellular reprograming approach that employed a Newtonian plasma fluid without incorporating the RBCs to plasma cultured protocol (Sharma et al., 2022) Further importance of co-developing hematovascular system in embryonic CSF development is reflected from the studies that support the dual role of neuroepithelium lining and blood components of adjoining vascularized niches in contributing to CSF composition (De Lange and Danhof, 2002, Wislocki GB et al.,1958). Prior to choroid plexus and blood-brain barrier development, growth factors such as fibroblast growth factor 2 and other blood-born peptides are also reported to freely enter the embryonic CSF via transcellular route that spans vascular endothelial cells in the ventral mesencephalon and the anteroventral part of the prosencephalon (Wislocki GB et al.,1958, Chau et al., 2015, Liddelow et al., 2013).

Brain vasculature is marked by earliest blood supply to the embryonic nervous system that comes from a perineural vascular plexus covering the surface of the neural tube around weeks 3 and 4 (Zhu et al., 2018, Knott et al., 1997), followed by relatively late emerging (around eighth gestational week) intrinsic vascular plexus that penetrates the brain parenchyma and is marked by the fusion of the basement membranes of the capillary and glia limitans externa (outer glial membrane of the brain made up of glial end feet) (Liddelow et al., 2014, Dziegielewska et al., 1988), suggestive of a close spatial-temporal developmental trajectories of endothelial and glial cells. The leptomeningeal surface vasculature is contributed by the surface plexus, whereas the intrinsic brain vasculature is contributed by mutual neuro-hematovascular signaling involving VEGF-A-VEGFR2 axis. The neuroectoderm secreted VEGF-A activates VEGFR2 receptors on the angioblasts of the perineuronal vascular plexus to initiate the formation of endothelial sprouts via sprouting angiogenesis that facilitate the formation of the intrinsic vascular plexus, which later penetrates the brain parenchyma. Neurons and glia are two important sources of VEGF-A during embryonic development, and the coordinated interaction between the cells of neurovascular unit drives vascular development during embryonic and postnatal life (D’Gama et al., 2021). Another important feature of developing neurovascular niches in brain is the induction of choroid plexus fate by repression of neuronal transcription factors in neuroepithelial cells in the rhombic lip region followed by vascularization of the choroid plexus via signaling between the roof plate neuroepithelium, choroid plexus epithelium, and pericytes that drives the formation of fenestrated capillaries in the choroid plexus (Abdelhamed et al., 2018) which again reflects the flexibility of transport of certain (large) molecules and electrolytes from blood plasma to the CSF and their direct delivery to the ventricles (Brightman and Reese, Brightman 1968, Doolin and Birge, Doolin and Birge, Vong et al.)Retinol-binding protein is reported to transport all-trans-retinol from the embryonic plasma to the embryonic CSF where it is associated with neuronal proliferation and differentiation (Keep and Jones, 1990, Quay). Embryonic CSF has a complex protein composition when compared to blood plasma, and it also differs from that of the adult CSF (Kuro-O* et al., 1997),Zhu et al., 2018, Knott et al., 1997, Dziegielewska et al., 1988, Liddelow et al., 2014. Early CSF that is contributed by neural tube trapped amniotic fluid and th e developing CSF proteomes diverge during development following the closure of anterior neuropore, and these relatively changing CSF protein compositions are linked to neural progenitor cell self-renewal (Liddelow, 2015) However, CSF protein composition changes during development are challenging to monitor due to anatomical constraints to sampling, nevertheless immunohistochemical staining for plasma proteins in choroid plexus cells demonstrate overlapping plasma protein profiles in subpopulations of choroid plexus epithelial cells. Importantly, the numbers of these plasma protein-containing cells are generally higher in choroid plexuses of brains from larger animals and the proportion of plasma protein-containing cells is higher earlier in development than in postnatal life (Jacobsen M et al 1982, Liddelow et al., 2010, (Knott et al., 1997)). Tightly controlled plasma protein transport in the developing choroid plexus is reported when compared with the adult, where there appears to be a loss of specificity and several plasma proteins can be transported by the same cell (Liddelow et al., 2013, Liddelow et al., 2014). This presence of plasma proteins suggests an integral role of protein transport from blood to the CSF via the interstitial space of the choroid plexuses and has been demonstrated across species (Knott et al., 1997, Dziegielewska et al., 1988, Liddelow et al., 2009). Interestingly the CSF turnover in the developing brain is slower as compared to the adult brain resulting in higher protein concentration (a feature of plasma, this suggested that transport results in a high concentration of individual plasma proteins and total protein concentration in CSF in the developing brain compared with the adult (Johansson et al., 2008, Dziegielewska et al., 1981, Knott et al., 1997, (Dziegielewska et al., 1988,Cavanagha et al., 1982). The co-evolution perspective of neuro-hematovascular niches is not just limited to developmental process, it is also reflected during ischemic stroke linked adult neurogenesis, where deregulated hemostasis process triggers neuronal remodeling. However, future lineage tracing studies will decipher if an *in vivo* common bipotential ecto-mesodermal population (reminiscent of ecto-mesodermal neural crest stem cells that contribute to peripheral nervous system) contributing to CNS neurogenesis exists in blood.

## Methodology

Culturing neuro-hematovascular niches from autologous plasma: A blood sample was obtained from a healthy subject who had no underlying health conditions at the time of blood withdrawal. Autologous blood components, specifically plasma, white blood cells (WBCs), and red blood cells (RBCs), were separated using density dependent centrifugation with Granluosep (HiMedia Granulosep 1119) and Hisep (HiMedia Hisep 1077). Buffy coat containing PBMC’s and granulocyte were isolated and pelleted. 1 ml of plasma and 100 µl of the cell mixture per well is plated. Fo obtaining the conditions C1, C2, C3 and C4, RBC were added in increasing volumes, where C2 (2 µl per 1ml of plasma) served as the baseline RBC concentration, C3 contains twice that of C2 and C4 has thrice the concentration tha C2.The cells were incubated for two weeks (14 days) in a CO2 incubator at 37°C with 5% CO2, and routine monitoring was conducted using bright field microscopy.

Immunofluorescence: Cells and tissue were fixed with 4% paraformaldehyde (PFA) followed by treatment by 0.3% Triton X-100 and blocked using a freshly prepared blocking solution (2.5% BSA). Cells were then incubated with primary antibodies at the following dilutions: c-Kit (rabbit, Cell Signaling Technology, 3074s, 1:300), Stargazin (mouse, Invitrogen, Iba-1 (rabbit, Invitrogen, MA5-29012, 1:300), Dopamine Beta Hydroxylase (DBH) (sheep, Invitrogen, PA3925, 1:300), Beta -3 Tubulin (mouse, Invitrogen, MA119187, 1:300), VEGFR2 (rabbit, Invitrogen, PA1-16613, 1:300), CD40 (donkey, Invitrogen, PA5-142944, 1:300), GFAP (mouse, Invitrogen, MA515086, 1:300), Neun (rabbit, Invitrogen, PA5-78639, 1:300), VGLUT2 (mouse, Invitrogen, MA527613, 1:300), Tissue Growth Factor-beta1 (TGF-β1) (rabbit, Bioss, Bs-0086R,1:300), COL1A1 (donkey, Santa Cruz Biotechnology, sc-25974, 1:300),, VEGFR3 (mouse, Invitrogen, 14-5988-82, 1:300), FoxA2 (rabbit, Invitrogen, 1:300,701698), LEF1/ TCF1 (rabbit, NeoBiotechnologies, 51176-RBM1-P1, 1:300), Phospho-FGFR2 (rabbit, Invitrogen, PA5-105880), Ki67 (rabbit, Invitrogen, PA5-19462, 1:200), FGF-2 (mouse, Santa Cruz Biotechnology, sc271847, 1:300), PDGFR-Alpha (mouse, Santa Cruz Biotechnology, sc-398206, 1:300),Sox-2 (mouse, Santa Cruz Biotechnology, sc-365823, 1:200) Oct4 (mouse, Santa Cruz Biotechnology, sc-5279), ADRA 2A (rabbit, Invitrogen, PA1-048,1:300), VEGFR3 (mouse, Invitrogen, 14-5988-82,1:300), AchR (rabbit, Invitrogen,YK4127703, 1:300). Secondary antibodies used were goat Alexa Fluor 488 and 568 and donkey Alexa Fluor 647 conjugates (Invitrogen, 1:200) and Alexa Fluor 647 Phalloidin (Invitrogen, 1:500). DAPI was used in 1:20 dilution for staining nuclei.

Scanning Electron Microscopy: To understand the morphology and development of the various types of cell being developed in our culture, Cells were fixed with 4% paraformaldehyde, dehydrated in graded ethanol, and subjected to critical point drying. Samples were coated with 10□□ mm gold particles and analyse using a JEOL IT300 (v1.5) SEM. Morphology was examined at 30 kV using secondary and backscattered electron detectors, achieving magnifications up to 300,000x.

Cation Exchange-HPLC (Hemoglobin typing): Analytical HPLC quantification of hemoglobin (Hb) tetramers and individual globin chains was performed using IE and RP columns on an HPLC system.The Bio-Rad VARIANT II system was used for Hb typing through ion-exchange high-performance liquid chromatography (HPLC). The required volume of cell cultured in plasma (typically 20–25 µL) taken into a vial and was lysed with demineralized water by centrifuging it at 2500 RPM for 6 minutes. The sample was analysed via the software interface which automatically draw the sample into the system. The Hb variants (e.g., HbZ, HbE, HbA, HbF, HbA2, and abnormal variants like HbS, HbC, etc.) were isolated using a programmed buffer gradient oh HB testing buffers. The separated Hb types were detected using a UV detector (typically at 415 nm). The chromatograms were monitored, displayed on the screen in real-time. Each peak corresponds to a specific Hb fraction or variant. The retention time and area under the curve (AUC) are used to identify and quantify Hb types.

LCMS/MS and Bioinformatic analysis: Protein samples (50 µg) were reduced with 5 mM TCEP, alkylated with 50 mM iodoacetamide, and digested with trypsin (1:50, enzyme:lysate) for 16 h at 37 °C. Peptides were cleaned with C18 cartridges, dried, and resuspended in buffer A (2% acetonitrile, 0.1% formic acid). Mass spectrometric analysis was performed on an Easy-nLC 1000 coupled to an Orbitrap Exploris using a 15 cm C18 column. Peptides were separated over a 110-min gradient (0–40% buffer B: 80% acetonitrile, 0.1% formic acid) and analyzed in MS1 (R = 60,000) and MS2 (top 20, R = 15,000). Data were processed using Proteome Discoverer (v2.5) against the UniProt Human database with trypsin/P specificity and a 1% FDR cutoff. Carbamidomethylation (C) was set as a fixed modification, and oxidation (M) and N-terminal acetylation as variable modifications. Pathway and network analysis were performed using STRING, DAVID, KEGG, ShinyGO, Enrichr, Cytoscape, and FunRich. Proteins with p < 0.05 and log2FC > 1 were analyzed for biological processes, molecular functions, and pathway interactions.

Thrombin generation chromogenic assay: The thrombin generation assay utilized as a chromogenic substrate (β-Ala-Gly-Arg p-nitroanilide diacetate) (Cat no #T3068) The substrate was reconstituted to a 5 mM stock and diluted to a working concentration of 0.5 mM. Thrombin standards (100 µg/ml stock) were prepared in a two-fold serial dilution (100–1.56 µg/ml), with PBS as a blank. For the assay, 60 µl of sample, standard, or blank was mixed with 75 µl of 50 mM Tris-HCl buffer (pH 7.4), and an initial reading was recorded at 405 nm. Following the addition of 15 µl of 5 mM substrate, the reaction was incubated at 37°C for 15 minutes, and a final reading was obtained. Results were calculated and ANOVA was performed on data from conditions C1–C4 (increasing RBC levels).

Plasmin Generation Assay: The assay utilized a chromogenic substrate (N-Tosylglycyl-L-prolyl-L-lysine 4-nitroanilide acetate, CAS No. 88793-79-7) cleaved by plasmin to release a fluorophore. A 78.8 mM stock substrate was prepared in ethanol and diluted to a working concentration of 200 µM. Thrombin standards (100 µg/ml stock) were prepared in a two-fold dilution series (100–1.56 µg/ml), with PBS as the blank. A 50 mM Tris-HCl buffer (pH 7.4) was used. For the assay, 50 µl of sample, standard, or blank was mixed with 49.75 µl of buffer, and an initial reading at 405 nm was taken. Following the addition of 0.25 µl of substrate, the reaction was incubated at 37°C for 30 minutes, and a final reading was recorded. Results were calculated and ANOVA was performed on data from different conditions.

MMP fluorogenic activity assay: MMP activity was measured using the fluorogenic substrate Mca-Lys-Pro-Leu-Gly-Leu-DNP-Dpa-Ala-AR-NH2 (CAS No. 720710-69-0). A 10 mM stock was prepared in DMSO and diluted to a working concentration of 5 µM in 25 mM Tris HCl buffer (pH 8) containing 80 mM CaCl□. Reactions were performed in 100 µL volumes, comprising 50 µL of sample or blank and 50 µL of substrate solution.The assay was conducted in black-walled 96-well plates (Nunc), and fluorescence was recorded using a multimode reader with excitation at 320 nm and emission at 405 nm, at 5-minute intervals for 30 minutes. ANOVA was performed on data from conditions C1–C4 (increasing RBC levels).

IGF-1 Quantitative Assay: The Elecsys IGF-1 assay, was performed on our cultured plasma using Cobas immunoassay analyzers, which quantitatively measures IGF-1 in human serum or plasma using a two-step electrochemiluminescence immunoassay (ECLIA). IGF-1 is released from IGF-binding proteins with an acidic buffer, then binds to a biotinylated monoclonal antibody. A ruthenium-labeled antibody binds to another epitope, forming a sandwich complex. Streptavidin-coated magnetic microparticles capture the complex, and unbound substances are washed away. The emitted light is measured, and IGF-1 concentrations are reported in ng/mL.

Statistical Analysis: We conducted statistical analyses on the following functional assays: Thrombin generation (n=10), Plasmin generation (n=10), and Matrix metalloproteinases (MMP) (n=20), with results displayed in Figures 4E, 4F, and 4G, respectively. Additionally, a functional assay on IGF-1 (n=7) is shown in Figure 3F. For the analyses of Thrombin generation, Plasmin generation, and MMP, we used ANOVA (non-parametric) for multiple comparison groups. For IGF-1, a non-parametric t-test was employed to compare two groups, utilizing GraphPad Prism 8.0. We defined statistical significance as p-values less than 0.05. Asterisks indicate the following p-value thresholds: p < 0.0001 = ****; p < 0.001 = ***; p < 0.01 = **; p < 0.05 = *; ns = not significant.

## Resource Availability

## Corresponding author

Further information and requests for resources, materials, or data should be directed to the corresponding author, **Maryada Sharma**

## Materials and availability

All antibodies, and other reagents used in this study are commercially available. Data and code availability This study did not generate new unique reagents. The raw proteomics data and confocal microscopy images generated during this study are available from the corresponding author upon reasonable request.

## Ethics Statement

The study has been approved by the institutional ethics committee with the following approvals: IEC-05/2020-1672 and PGI-IC-SCR-107-2020-1418.

## Acknowledgements

We acknowledge SERB media cell for featuring our work on its official portal https://dst.gov.in/new-prototype-developed-generate-neurovascular-tissuesorganoidsautologous-blood-can-help-precision; and constant funding support-SERBSPR/2019/1447; SERB-CRG/2019/6755; SERB-CRG/2022/4946).

## Author contributions

Conceptualization-M.S.

Data curation-R.A., A.B., P.P.

Formal analysis -R.A.

Funding acquisition – M.S.

Investigation – R.A.

Methodology –M.S.

Project administration-M.S., R.D., S.S., T.G.

Resources – M.S., N.K., J.B., G.N., R.V., S.K., S.S., R.D., T.G., S.B., A.P., R.P., M.L., S.R., S.C.,S.A

Supervision-M.S., R.D., S.S.,T.G.

Validation – R.A., M.S.

Visualization – R.A., A.B.

Writing – original draft-A.B., R.A., M.S.

Writing – review & editing-A.B., S.S., M.S.

R.A.-Rhythm Arora

A.B.-Alka Bhardwaj

M.S.-Maryada Sharma

N.P.-Naresh K Panda

J.B.-Jaimanti Bakshi

R.V.-Ramandeep Singh Virk

R.D.-Reena Das

S.S-Sanhita Sinharay

P.P.-Pooja Patkulkar

T.G.-Tulika Gupta

S.B.-Sanjay Kumar Bhadada

A.P.-Arnab Pal

R.P.-Rimesh Pal

M.L.-Meenakshi Pal

G.N.-Gyanaranjan Nayak

S.K.-Sourabh K Patro

S.C.-Seema Chhabra

S.R.-Sajid Rashid

S.A.-Sanjeev Chhabra

